# Computer-assisted EEG diagnostic review for idiopathic generalized epilepsy

**DOI:** 10.1101/682112

**Authors:** Shannon Clarke, Pip Karoly, Ewan Nurse, Udaya Seneviratne, Janelle Taylor, Rory Knight-Sadler, Robert Kerr, Braden Moore, Patrick Hennessy, Dulini Mendis, Claire Lim, Jake Miles, Mark Cook, Dean Freestone, Wendyl D’Souza

## Abstract

Epilepsy diagnosis can be costly, time-consuming and not uncommonly inaccurate. The reference standard diagnostic monitoring is continuous video-EEG monitoring, ideally capturing all events or concordant interictal discharges. Automating EEG data review would save time and resources, thus enabling more people to receive reference standard monitoring and also potentially herald a more quantitative approach to therapeutic outcomes. There is substantial research into automated detection of seizures and epileptic activity from EEG. However, automated detection software is not widely used in the clinic; and, despite numerous published algorithms, few methods have regulatory approval for detecting epileptic activity from EEG.

This study reports on a deep learning algorithm for computer-assisted EEG review. Deep, convolutional neural networks were trained to detect epileptic discharges using a pre-existing dataset of over 6000 labelled events in a cohort of 103 patients with idiopathic generalized epilepsy (IGE). Patients underwent 24-hour ambulatory outpatient EEG, and all data was curated and confirmed independently by two epilepsy specialists (Seneviratne et al, 2016). The resulting automated detection algorithm was then used to review diagnostic scalp-EEG for seven patients (four with IGE and three with events mimicking seizures) to validate performance in a clinical setting.

The automated detection algorithm showed state-of-the-art performance for detecting epileptic activity from clinical EEG, with mean sensitivity of >95% and corresponding mean false positive rate of 1 detection per minute. Importantly, diagnostic case studies showed that the automated detection algorithm reduced human review time by 80%-99%, without compromising event detection or diagnostic accuracy. The presented results demonstrate that computer-assisted review can increase the speed and accuracy of EEG assessment and has the potential to greatly improve therapeutic outcomes.

## 1. Introduction

Epilepsy is a severe neurological disease typified by recurrent seizures. The underlying causes of epileptic seizures are varied and include structural abnormalities, genetic mutations, and many other unknown mechanisms [1]. This heterogeneity makes epilepsy diagnosis challenging. The widely accepted operational definition requires the greater tendency to recurrent seizures [2]. However, other conditions can also provoke epileptic seizures, including metabolic derangements or intoxication. Furthermore, fainting, psychogenic causes, or sleep disturbances may frequently present with symptoms mimicking seizures. Consequently, epilepsy diagnosis can involve a complex and costly process with a rate of misdiagnosis ranging from 30%-70%, depending on the type of testing conducted [3]. Although, epilepsy remains a clinical diagnosis, for research and higher clinical precision the diagnostic reference standard is the capture of typical events during simultaneous video-EEG monitoring. However this typically takes several days and sometimes up to one week [4]. This testing generates vast amounts of data that must be reviewed in order to determine whether the EEG signatures of an epilepsy syndrome are present. Data recorded during a clinical seizure is the most valuable test; however, seizures are relatively infrequent and diagnostic EEG must still be thoroughly reviewed for more subtle interictal epileptic activity such as spike-wave discharges, bursts, focal slowing, or other diagnostic markers. Automating diagnostic reviewing has the potential to save time, reduce costs and improve accuracy, thus enabling more people to receive reference-standard monitoring and improving diagnostic outcomes in epilepsy.

There has been decades of research into automated detection of seizures and epileptic activity from EEG, with hundreds of papers published each year [5–7]. The advent of machine learning and artificial intelligence has sparked a new wave of interest in developing generalizable algorithms that can be trained to recognise epileptic activity in EEG data. The seizure detection problem has even reached mainstream data science, with recent Kaggle competitions attracting thousands of entrants whose submissions covered a range of machine learning methods from deep convolutional neural networks to extreme gradient boosting [8–10]. A recent review of seizure detection from scalp EEG reported good performance from available algorithms, with sensitivities between 75% and 90% and false positive rates of between 0.1 and 5 per hour [6]. However, a large study of 1,478 ambulatory EEG studies found that commercially available automated detectors correctly identified electrographic seizures in only 53% of cases [11]. For diagnostic review, it is arguably as important to detect interictal epileptiform discharges (IEDs) in addition to seizures, which increases the difficulty of algorithmic detection, as there is often little consensus from human reviewers on what constitutes an IED [12–14]. Automated IED detectors have been developed over decades [15–17], often with a focus on clear morphological features, such as “spikes”. A huge array of features and classifiers have been employed in automated spike detection from EEG, with reported sensitivities ranging from 60%-100% and false positive rates of 0.1 to 6 per minute [7]. Nevertheless, despite thousands of published spike detection algorithms with excellent performance, there remains just one FDA approved method for detecting epileptic activity from EEG [14].

There are several hurdles to translating automated IED detectors from a research to a clinical setting. Clinical recordings may vary considerably between different epilepsy centres [18]. Scalp EEG used for diagnostic review also contains many artefacts, which may be exacerbated in a home monitoring setting. Seizures and IED morphology can be highly varied between individuals with different epilepsy syndromes [19]; yet, for practical utility, automated review algorithms must be able to generalize to a broad patient population [20]. Because of these challenges, it is generally accepted that automated detection algorithms are not yet able to replace human reviewers. A common approach in clinical research is computer-assisted review, where an algorithm first analyses all the data and flags suspect activity. The computer detections are then checked by an expert human reviewer. This approach enables an achievable balance between high sensitivity while minimising false positive rates. High sensitivity is important to quantify the frequency of IEDs, for instance as a potential measure of therapeutic efficacy or disease burden. In other cases, lower false positive rates may be more important, such as for diagnostic confirmation, or source localisation for surgical planning. However, false positive rates should be low enough to ensure that human reviewers do not have to re-check large amounts of data, rendering computer assistance redundant.

Machine learning, and in particular deep learning, has rapidly become a useful technology for a wide variety of applications, including interpretation of medical diagnostic data sources [21–23]. Deep learning uses labelled data sets to generate highly-complex models (“neural networks”) that can classify new data into discrete categories. Deep learning may be particularly useful for automated IED detection, because neural networks can learn to recognise generalisable signal features from a diverse training examples, such as EEG from different patients. Neural networks have been used for EEG spike detection for decades [24]; however, deep learning performance has not yet surpassed more traditional detection algorithms based on pre-defined signal features [7,25]. One potential hurdle is that deep learning requires large amounts of well-curated data compared to other machine learning techniques, due to the high number of model parameters. Hence, large numbers of digitized EEG records, with corresponding annotations, are required to train neural networks to detect epileptiform events. Recent technological advances, including disposable electrodes, improved battery life and wireless data transmission, have made it easier to record and save large databases of EEG [26], leading to the ability to rigorously evaluate deep learning algorithms for IED detection in a clinical setting.

## 2. Material and Methods

This study presents a deep learning algorithm for computer-assisted EEG review. Here, we report on results from a cohort of patients with idiopathic generalized epilepsy (IGE). Clinical case studies are also presented to demonstrate the difference in computer-assisted review between patients with a prior diagnosis of IGE and patients without epilepsy. This study was conducted with approval from the Human Research Ethics Committees of Monash Health and St. Vincent’s Hospital, Melbourne. All subjects provided written informed consent in keeping with the Declaration of Helsinki.

### 2.1 Data

Two datasets were used in this study, an evaluation dataset of 103 patients was used to both train neural networks and test performance (using five-fold cross-validation, described in detail in subsequent sections). In addition, to testing performance with this large cohort, a smaller cohort of seven patients with either IGE or seizure mimickers was used to provide a proof-of-concept for clinical utility. Further detail on the data, neural network development, and performance assessment is provided in the following sections.

#### 2.1.1 Evaluation dataset

Algorithm evaluation was based on 24-hour ambulatory EEG records from 103 patients diagnosed with IGE [27–29]. For a detailed description of the cohort, see [29]. Briefly, 24-hour ambulatory EEG recordings were undertaken using a standard 10-20 protocol at a single tertiary hospital in Melbourne, Australia (St. Vincent’s Hospital Melbourne). EEG was acquired using a 32 channel Siesta EEG system (Compumedics Ltd., Melbourne, Australia). Anti-epileptic drugs were not changed during the recording. The patient cohort had a mean age of 28 ± 10.7 (range 13-58) and was 66.7% female. Further clinical characteristics are available in [27]. The complete dataset was reviewed by a human epileptologist and IEDs were marked. In total, there were 6983 labelled IEDs (range of 0 to 556 IEDs per patient, mean 67.8 IEDs per patient)

#### 2.1.2 Clinical case studies

Clinical case studies included seven patients (four with IGE and three without epilepsy) undergoing at-home ambulatory video-EEG monitoring. Case studies provided a proof-of-concept for the classification algorithm performance in a diagnostic clinical setting. Data was recorded for one day to approximately one week using silver/silver chloride EEG scalp electrodes with a standard 10-20 configuration. Recording and human review was undertaken by Seer, a home monitoring service for epilepsy diagnosis. Deidentified patient EEG recordings were obtained for the purposes of this study, as well as the final clinical diagnosis from the treating neurologist. Clinical EEG was automatically labelled by the detection algorithm, and all labels were reviewed by initially by experienced neurophysiologists (authors JT, RKS) and subsequently by an epilepsy specialist (MC). At the time of EEG monitoring, Patients 1-4 already had a confirmed diagnosis of IGE. Patients 5, 6, and 7 had no diagnosis and were undergoing diagnostic monitoring. Following diagnostic monitoring, Patients 5 – 7 were all diagnosed with PNES by the referring neurologist. Table 1 provides a summary of patient demographics.

**Table 1.** Patient demographics for clinical case studies.

| Patient | Diagnosis | Recording duration (hours) | Age | Gender |
| --- | --- | --- | --- | --- |
| 1 | IGE | 21 | 34 | F |
| 2 | IGE | 24 | 26 | F |
| 3 | IGE | 21 | 22 | M |
| 4 | IGE | 95 | 59 | M |
| 5 | None (PNES) | 166 | 30 | F |
| 6 | None (PNES) | 155 | 29 | M |
| 7 | None (PNES) | 168 | 13 | M |
| | | Mean ± SD: 92.9 ± 70.6 | Mean ± SD: 30.4 ± 14.3 | Female – 43% |

### 2.2 Neural networks

Figure 1 shows a schematic of deep neural network, where segments of EEG data are used as inputs and the network is trained to output a certainty (between 0 and 1) that the input data is epileptiform. The neural network was developed using Pytorch version 1.01 [30] and Python version 3.6.

**Figure 1.**
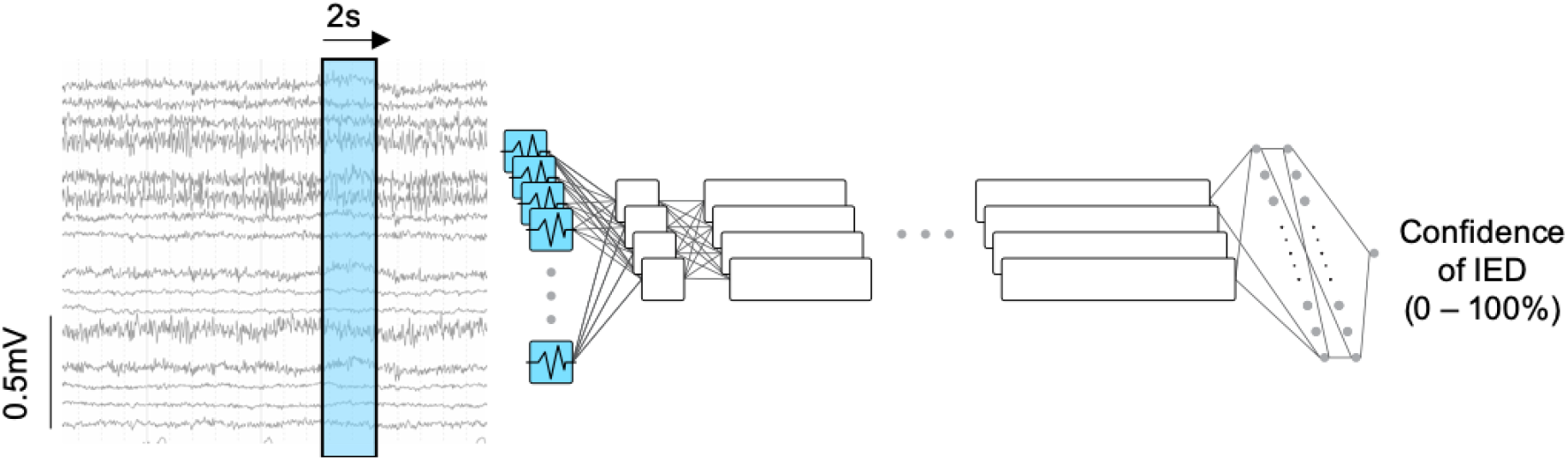
Neural network training. A 2s window of scalp EEG data is used as a training input for deep neural networks. Training data is input in batches into the first convolution layer of the neural network. Following several layers of convolutional filters and fully-connected neurons the final binary neuron layer outputs a confidence level (0-100%) that each EEG window is epileptic. The weights between network neurons are learned through repeated presentation of training examples.

#### 2.2.1 Network structure

Deep neural networks are computational models inspired by the architecture of the brain. Neural networks consist of many layers of interconnected “neurons” that process information via basic non-linear operations (i.e. step- or sigmoidal functions). Convolutional neural networks are a class of neural network that also include layers of 2D filters that are convolved with inputs. These filter layers are particularly useful to classify higher dimensional data, such as images or multi-channel EEG [23,31]. During learning, the connections (“weights”) between layers are updated through repeated presentation of training examples and their corresponding labels (in this case, “epileptic” and “non-epileptic”). In this way, the neural network “learns” to represent highly abstract features of the EEG through millions of non-linear combinations of filter and neural operations. This high-dimensional, abstract representation of EEG allows neural networks to efficiently decode information, in order to differentiate between signal classes of interest, or detect anomalies in data. In this study, a neural network was trained to recognise IEDs using training data from the evaluation cohort of 103 patients. The neural network consisted of seven convolutional (filtering) layers followed by two fully connected (neuron) layers. Convolutions were applied across the time dimension only, to build up time-based EEG features in each channel individually. Table 2 shows the size (number of filters or neurons) and order of the neural network layers.

**Table 2.** Neural network layer structure. A summary of the layers (type and size) used for neural network models. See Supplementary Table 1 for full network parameters.

| Layer | Type | Size |
| --- | --- | --- |
| 1 | Convolution | 32 |
| 2 | Convolution | 64 |
| 3 | Convolution | 128 |
| 4 | Convolution | 256 |
| 5 | Convolution | 256 |
| 6 | Convolution | 256 |
| 7 | Convolution | 128 |
| 8 | Fully connected | 128 |
| 9 | Fully connected | 128 |
| 10 | Output | 1 |

#### 2.2.2 Network training and validation

The network was trained on 2s windows of EEG data. The data were passed through three Butterworth filters; a second order low-pass at 70Hz, a second order high-pass at 0.5Hz and a first order notch filter between 45 and 55Hz. The data were then normalised to have a standard deviation approximately equal to 1. The entire cohort of 103 patients contained over 6000 labels and a combined recording duration of over 100 days [29]. Stochastic optimisation [32] with a learning rate of 1e-3 and batch size of 2560 was used to train the network. Training was stopped after 800 batches to avoid overfitting. Each batch contained 50% positive samples (epileptiform EEG) and 50% negative samples (non-epileptic EEG). The network took five hours to train on an NVIDIA Tesla M60 GPU.

### 2.3 Performance evaluation

For the evaluation cohort of 103 patients, five-fold cross validation was used to validate algorithm performance. For each fold of the validation, 80% of the patients were used to train the neural network and the remaining 20% were used for testing. Testing was repeated five times with non-overlapping patient subsets. Patient data were not separated across training and testing subsets to avoid the algorithm being tested on data from the same patients used for training. Positive event detections were considered to be data windows that the algorithm labelled as epileptic with at least 50% confidence.

Performance was measured using sensitivity and false positive rate per minute, which are standard metrics for IED detection algorithms [6]. Sensitivity, S, is given as:

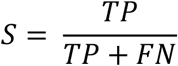

where TP are true positives (IEDs labelled by the algorithm and confirmed by specialist reviewer) and FN are false negatives (IEDs labelled by specialist reviewer but missed by the algorithm). It is only possible to detect false negatives if an expert human reviewer inspects and annotates the entire dataset, to provide a ground truth. In clinical case studies, only the events flagged by the algorithm were checked by human reviewers. Therefore, for clinical case studies we report on precision rather than sensitivity. Precision, P, is also referred to as the positive predictive value, and is given as the ratio of true positives over the total number of labels flagged by the algorithm:

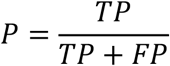

where FP are false positives. False positives are labels that were marked by the algorithm but not by human reviewers. The false positive rate is given as the number of false positives per minute.

In addition to sensitivity and false positive rate, we quantified the amount of remaining data that must be reviewed by a specialist following computer detection. These “data views” were used because, in a clinical setting, human reviewers require a larger window of data around a suspect IED to effectively determine whether the identified event is epileptiform or benign. For the clinical case studies, a 10s viewing window of EEG was used to determine whether suspected IEDs were real (true positives) or false alarms. Therefore, 10s was used as the default view size to compute data reduction. In practice, the view size may change depending on the type of event (seizure vs IED, focal vs generalised) or type of epilepsy. The number of data views per patient depends on the false positive rate as well as the distribution of false positives. For instance, if false positives are evenly dispersed throughout the data then the number of data views is the same as the number of false positives. However, if false positives tend to be clustered within the same 10s window, which often occurs in periods of noise or movement, then the number of data views is less than the number of false positives. In other words, knowing just the false positive rate does not necessarily provide a good indication of an algorithm’s clinical utility. Even a relatively high false positive rate can provide significant data reduction, leading to valuable time savings for clinicians. In this study we report the number of false positives as well as the number of data views per minute. The data views were then used to calculate clinical data reduction, which is the percentage of data remaining that still requires human review (i.e. the number of 10s data views divided by the length of the data).

## 3 Results

### 3.1 Neural network performance

Neural networks were evaluated on 103 patients with over 6000 labelled IEDs and over 100 days of recording (mean recording duration of 25 hours per patient). Performance was calculated using IED detections where the algorithm output confidence levels of 50% or greater. Over the five folds of cross validation the mean sensitivity was 96.7% (95% confidence interval using the standard error of the mean, CI [95.0, 98.3]). The mean false positive rate was 1.16 per minute (CI [1.00, 1.32]). The mean data reduction was 84.2% (CI [82.2 86.3]).

Figure 2 shows a plot of mean sensitivity and false positive rates as the detection confidence level was varied between 50% to 99%. Higher confidence levels result in fewer false positives, at the expense of missing more IEDs. At a confidence level of 99%, the mean false positive rate was 0.14 per minute (CI [0.10 0.17]) and sensitivity was 85.0% (CI [80.9 89.1]).

**Figure 2.**
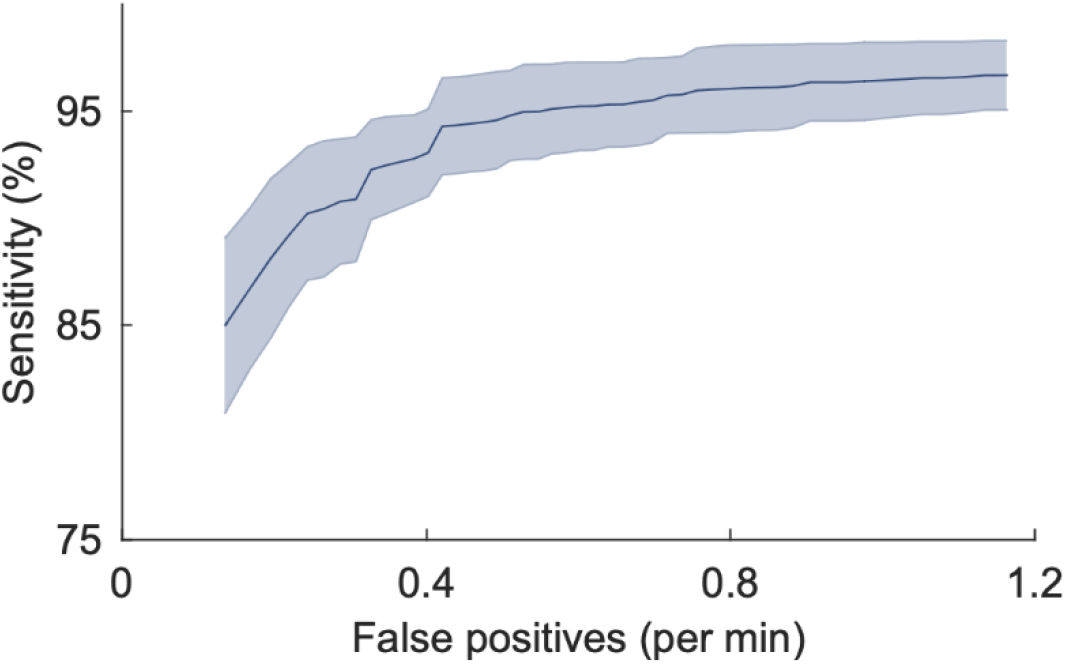
Sensitivity vs false positive rate for different confidence levels. Performance was based on IED detection for the 103 patients in the validation cohort. The plot shows mean false positive rates (x-axis) compared to sensitivity (y-axis). Confidence levels were varied from 99% down to 50% (left to right). Mean values were computed over the five subsets of cross validation. The shaded error bar shows the 95% confidence interval calculated from the standard error of the mean.

Figures 3 and 4 show the distributions of false positive rates and data reduction for each validation subset (where a subset consists of 20% of the patients). It can be seen that the false positive rate was generally between 1 and 2 events per minute, leading to data reduction of over 80%.

**Figure 3.**
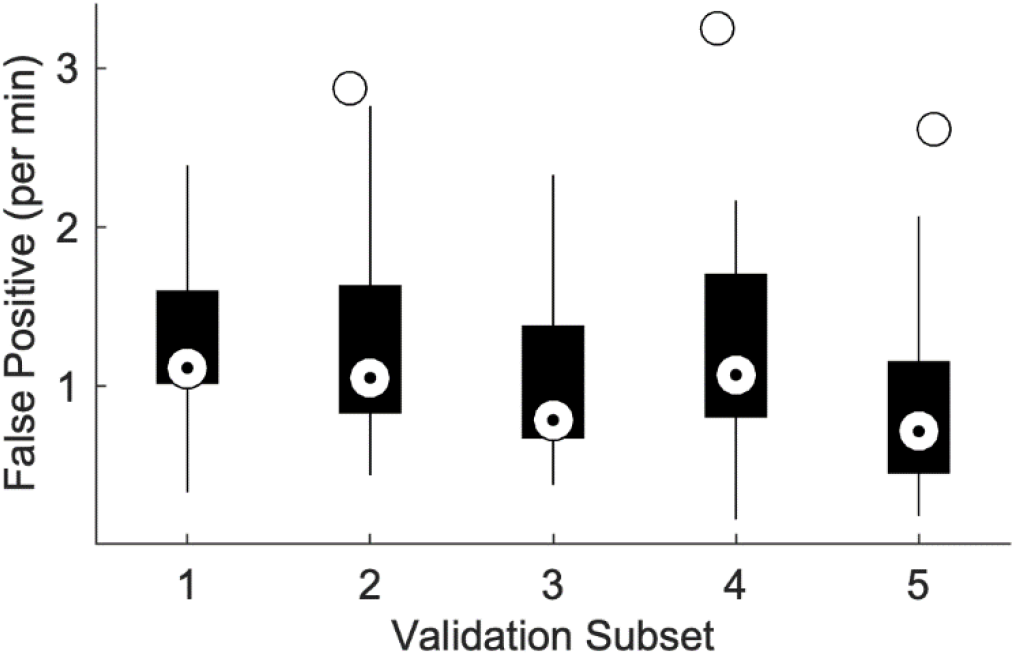
False postive rate distribution. Box plots show the distribution of false positive rates (y-axis) for each of the validation subsets (x-axis). Validation subsets contained 20% of the patient cohort (non-overlapping). Upper and lower box edges represent the 75^th^ and 25^th^ percentiles respectively. Upper and lower whiskers represent the 95^th^ and 5^th^ percentiles. White box markers show the median. Outliers are shown as black circles.

**Figure 4.**
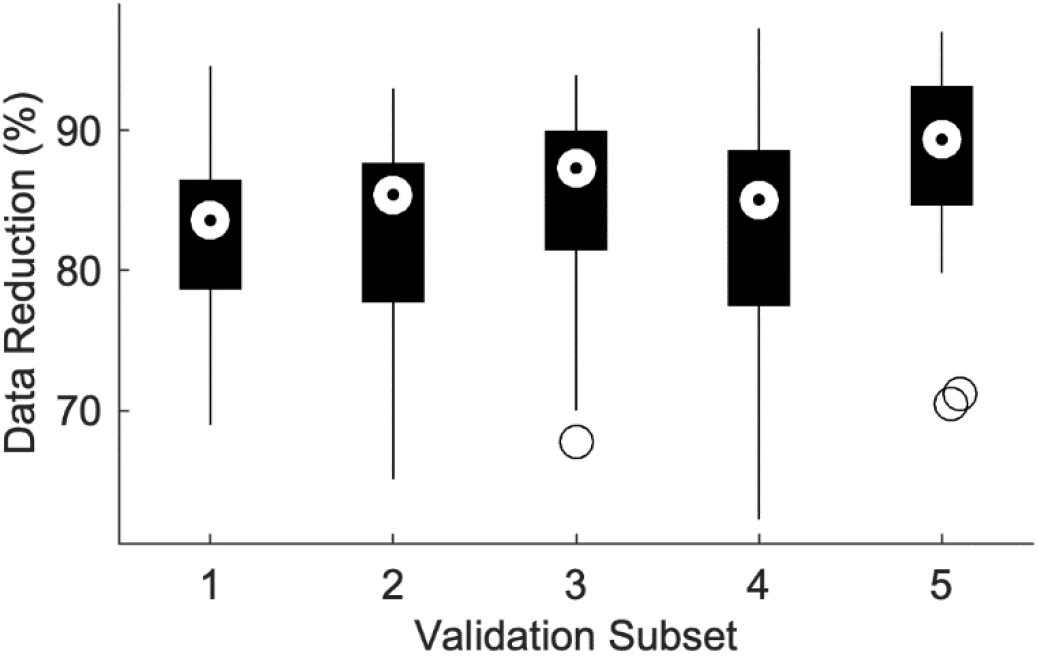
Data reduction distribution. Box plots show the distribution of data reduction (y-axis) for each of the validation subsets (x-axis). Validation subsets contained 20% of the patient cohort (non-overlapping). Upper and lower box edges represent the 75^th^ and 25^th^ percentiles respectively. Upper and lower whiskers represent the 95^th^ and 5^th^ percentiles. White box markers show the median. Outliers are shown as black circles.

### 3.2 Clinical case studies

Table 3 summarises the result of clinical case studies. It can be seen that the false positive rates were within the range of those seen in the validation dataset (Figure 3). Sensitivity was not assessed in the clinical case studies because only events labelled by the algorithm were reviewed by trained neurophysiologists; therefore, it is possible that some IEDs were missed (false negatives). Precision (the proportion of detected events that were true positives) was generally low. Patients 5, 6, and 7 had no epileptic events and were ultimately diagnosed with PNES. All patients had less data views than false positives, indicating some clustering of their false positive detections. The data reduction ranged from 65% to 90% over all patients and was greater than 80% for the patients without epilepsy.

**Table 3.**
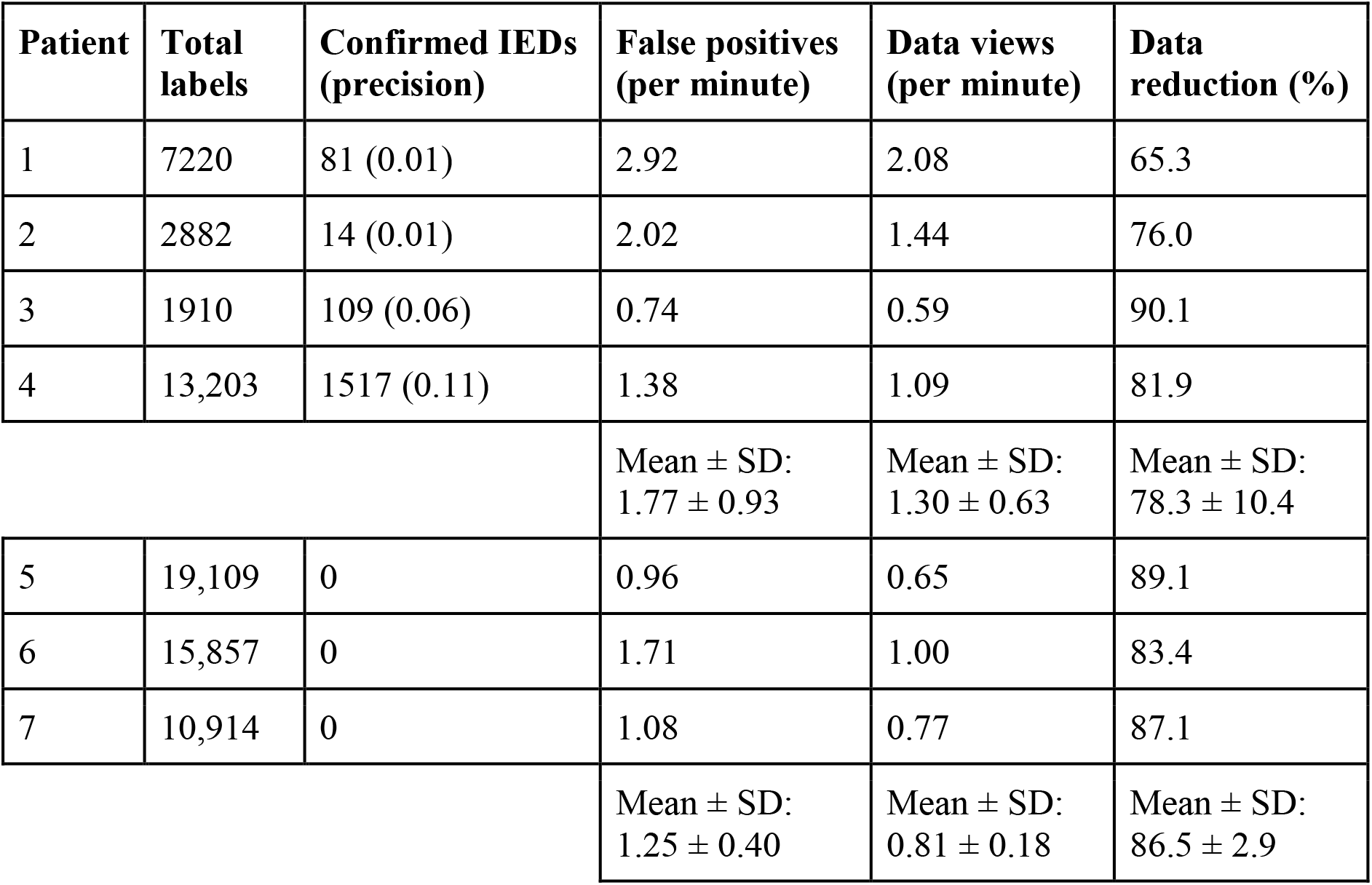
Performance results for clinical case studies. Data was labelled as epileptic if the neural network output >50% confidence of an IED. The number of total detections and confirmed epileptic events are reported with the associated precision. The rate of false positives and data views per minute are reported as well as the corresponding amount of data reduction. Summary statistics were computed separately for patients with IGE (Patients 1-4) and patients with PNES (Patients 5-7).

| Patient | Total labels | Confirmed IEDs (precision) | False positives (per minute) | Data views (per minute) | Data reduction (%) |
| --- | --- | --- | --- | --- | --- |
| 1 | 7220 | 81 (0.01) | 2.92 | 2.08 | 65.3 |
| 2 | 2882 | 14 (0.01) | 2.02 | 1.44 | 76.0 |
| 3 | 1910 | 109 (0.06) | 0.74 | 0.59 | 90.1 |
| 4 | 13,203 | 1517 (0.11) | 1.38 | 1.09 | 81.9 |
| | | | Mean ± SD:<br>1.77 ± 0.93 | Mean ± SD:<br>1.30 ± 0.63 | Mean ± SD:<br>78.3 ± 10.4 |
| 5 | 19,109 | 0 | 0.96 | 0.65 | 89.1 |
| 6 | 15,857 | 0 | 1.71 | 1.00 | 83.4 |
| 7 | 10,914 | 0 | 1.08 | 0.77 | 87.1 |
| | | | Mean ± SD:<br>1.25 ± 0.40 | Mean ± SD:<br>0.81 ± 0.18 | Mean ± SD:<br>86.5 ± 2.9 |

In some use-cases it may be more desirable to achieve higher precision (more data reduction) at the expense of reduced detection sensitivity. Because neural networks output the confidence of IEDs, the detection threshold can easily be tuned to achieve different performance specifications. Therefore, in addition to the standard detection performance shown in Table 3, we evaluated the “high precision” output of the algorithm, where IEDs were only labelled when the neural network had greater than 99% confidence (compared to 50% confidence using in Table 3). Table 4 shows detection performance using the high precision threshold. In Table 4, it was also possible to calculate detection sensitivity as the ratio of true positive detections relative to all IEDs identified following specialist review (i.e. the confirmed IEDs in Table 3). From Table 4 it can be seen that the high precision detection enables greatly increased data reduction (more than 85% for IGE patients, and more than 99% for non-epileptic patients). IED detection remained accurate, with sensitivities ranging from 78% to 99% (note that non-epileptic patients had no confirmed IEDs).

**Table 4.** High confidence performance for clinical case studies. Data was labelled as epileptic if the neural network output >99% confidence of an IED. The number of labels and confirmed epileptic labels are reported along with the sensitivity (compared to the true labels marked in Table 3). The rate of false positives and data views per minute are reported as well as the corresponding amount of data reduction. Summary statistics were computed separately for patients with IGE (Patients 1-4) and patients with PNES (Patients 5-7).

| Patient | Total labels | Sensitivity (%) | False positives (per minute) | Data views (per minute) | Data reduction (%) |
| --- | --- | --- | --- | --- | --- |
| 1 | 1649 | 98.8 | 0.66 | 0.64 | 89.4 |
| 2 | 1565 | 92.9 | 1.09 | 0.80 | 86.7 |
| 3 | 97 | 86.7 | 0.02 | 0.04 | 99.4 |
| 4 | 3088 | 78.1 | 0.13 | 0.22 | 96.4 |
| | | | Mean ± SD: 0.48 ± 0.50 | Mean ± SD: 0.43 ± 0.35 | Mean ± SD: 93.0 ± 5.9 |
| 5 | 760 | N/A | 0.04 | 0.04 | 99.4 |
| 6 | 421 | N/A | 0.04 | 0.04 | 99.3 |
| 7 | 272 | N/A | 0.03 | 0.03 | 99.6 |
| | | | Mean ± SD: 0.04 ± 0.01 | Mean ± SD: 0.04 ± 0.01 | Mean ± SD: 99.4 ± 0.2 |

## 4 Discussion

Deep learning neural networks can be trained to accurately detect IEDs with high sensitivity, and false positive rates in patients with IGE compared to patients with PNES that are practical for retrospective analysis. Using a validation dataset of over 100 days continuous scalp-EEG recording in 103 patients, sensitivity was over 95%, with a false positive rate of approximately 1 detection per minute. In comparison, the only other commercially available IED detector for clinical use reports sensitivity of approximately 38% at the same false positive rate [14]. Although, it is important to note that our study was focused on IED detection for patients with IGE and cannot be directly extrapolated to IED detection performance evaluated against all types of epilepsy. On the other hand, patients with IGE have a particular need for accurate, automated IED detection from EEG, as the cumulative number and duration of their epileptic discharges is considered extremely important to quantify the burden of epilepsy [27,33]. To establish a diagnosis, people suspected of having IGE may undergo serial EEG monitoring to confirm a diagnosis, particularly when sleep is not captured [34,35], assess disease activity, including response to treatment, and subclinical events in relation to fitness to drive [36]. Reducing review time through automated event detection can facilitate more regular EEG monitoring and has the potential to greatly enhance epilepsy management for IGE cases.

The presented results also showed that deep learning is practical in a real diagnostic setting. Seven clinical case studies showed similar performance to the results using the validation cohort. It was difficult to evaluate detection performance for clinical case studies, as there was a small cohort and EEG recordings were not comprehensively reviewed by specialists. However, we note that false positive rates were similar compared to the validation dataset. For diagnostic purposes, automated detection should be designed to not miss any IEDs, hence relatively low precision was observed. Nevertheless, human review time for the clinical case studies was decreased by 65-90% using computer-assisted review (Table 3). In applications that require higher precision, neural network detectors can easily be set to reduce data further, simply by increasing the confidence threshold required for event detection. High precision detection results (Table 4) showed data reduction was greater than 85% for IGE patients, with sensitivities between 78% and 99%. For non-epileptic patients, high precision review gave data reduction of greater than 99%. Overall, automated detection enabled clinicians to review data between 5 and 100 times faster, underscoring the potential of computer-aided review to improve diagnostic efficiency. Technologies for using deep learning in a real-time, low-power context are emerging, which may allow for the use of deep learning in seizure alert systems [37].

This study presented results for a cohort of epilepsy patients with idiopathic generalised epilepsy; however, given appropriate training data, the same deep learning approach is capable of generalizing to a much wider patient cohort. Similarly, neural networks are capable of distinguishing multiple output classes [23], enabling an algorithm to automatically label discrete event types such as seizures, spikes, polyspikes, or other epileptiform activity. For some diagnostic cases it may be sufficient to only detect electrographic seizures rather than all IEDs. Although, there is a growing recognition of the value in measuring all epileptic activity in addition to simply counting seizures [27,33,38]. In any case, deep learning can address these different use cases, simply by training neural networks on labelled data relevant to the required application.

In addition to generalizing to other diagnostic cases, this deep learning algorithm could also be applied to detect IEDs in other applications that continuously record EEG, such as implantable devices. The development of EEG electrodes that can be worn continuously for longer periods [39], as well as the recent advent of minimally invasive sub-scalp recording devices [40,41] will create a growing demand for automated data review to reliably detect epileptic activity from EEG. At the same time, such devices will ensure an exponentially increasing amount of data becomes available to train deep learning algorithms for IED detection.

This study focused on diagnostic review for idiopathic generalised epilepsy, which is typified by morphologically characteristic IEDs known as generalised spike-waves and polyspike waves [35,42]. Not all epilepsy syndromes are characterised by distinct morphological EEG features; therefore, IGE represents a somewhat simplified use-case for automated IED detection compared to general epilepsy diagnosis. Future work will focus on validation of the deep learning algorithm for diverse epilepsy syndromes across a wider patient population. It is also important to note that IGE represents 20-30% of all epilepsies [43], making it an important use case for automated IED detection.

Another limitation when evaluating automated EEG review is the difficulty in obtaining reference-standard labels for IEDs. Multiple specialist reviewers often do not agree on whether an EEG waveform is epileptiform or not [13,14], which makes it difficult to judge a computer’s accuracy. In fact, agreement between reviewers decreases continually as more reviewers are asked to inspect the same EEG, suggesting that there is no exact consensus on what constitutes an epileptiform discharge [13]. Ultimately, the best test of performance of automated detection algorithms is how useful they are in a clinical setting, and automated review software should provide enough flexibility to support different clinicians’ decision-making [13]. Additionally, it is important to continue to increase the amount of standardised, large databases of reference labels available for development and testing of automated IED detection software.

## 5 Conclusion

This study implemented a deep learning algorithm to train neural networks to automatically detect epileptic events from scalp EEG of patients with IGE. The results demonstrated that automated detections can aid clinical practice by reducing the time required to review diagnostic studies, without compromising the accuracy of event detection. Ultimately, it is our hope that automated IED detection can improve the speed and accuracy of EEG review for a range of epilepsy syndromes. Automated data review has applications in diagnosis, ongoing management and implantable devices, with the potential to improve diagnostic and treatment outcomes for people epilepsy.

---

**Supplementary Table 1:** Table of neural network parameters. Each convolutional layer had a 1-D kernel size of 3 and a stride of 2 and was followed by a rectified linear unit (ReLU) function activation layer. The two fully connected layers of size 128 were also followed by a ReLU activation layer. The output layer used a sigmoid activation function to constrain the output between 0 and 1. Conv: Convolution

| Layer | Input<br>512 x 21 | Filters | Kernel<br>size | Stride | Dilation | Padding | Input<br>size | Output<br>size time<br>dim | Receptive<br>field time<br>dim | Output size<br>(EEG<br>channels) | Output<br>shape |
| --- | --- | --- | --- | --- | --- | --- | --- | --- | --- | --- | --- |
| 1 | Conv-1-32 | 32 | 3 | 2 | 1 | 0 | 512 | 255 | 3 | 21 | Batch x 32<br>x 255 x 21 |
| 2 | Conv-2-64 | 64 | 3 | 2 | 1 | 0 | 255 | 127 | 7 | 21 | Batch x 64<br>x 127 x 21 |
| 3 | Conv-3-128 | 128 | 3 | 2 | 1 | 0 | 127 | 63 | 15 | 21 | Batch x<br>128 x 63 x<br>21 |
| 4 | Conv-4-256 | 256 | 3 | 2 | 1 | 0 | 63 | 31 | 31 | 21 | Batch x<br>256 x 31 x<br>21 |
| 5 | Conv-5-256 | 256 | 3 | 2 | 1 | 0 | 31 | 15 | 63 | 21 | Batch x<br>256 x 15 x<br>21 |
| 6 | Conv-6-256 | 256 | 3 | 2 | 1 | 0 | 15 | 7 | 127 | 21 | Batch x<br>256 x 7 x<br>21 |
| 7 | Conv-7-256 | 128 | 3 | 1 | 1 | 0 | 7 | 5 | 255 | 21 | Batch x<br>128 x 5 x<br>21 |
| 8 | Dense-1-128 | 128 |  |  |  |  |  | 128 |  |  | Batch x<br>128 |
| 9 | Dense-2-128 | 128 |  |  |  |  |  | 128 |  |  | Batch x<br>128 |
| 10 | Dense-3-1 | 1 |  |  |  |  |  | 1 |  |  | Batch x 1 |

